# Skin DNA Methylation Encodes Multidimensional Facial Aging Phenotypes with Distinct Biological Architectures

**DOI:** 10.64898/2026.03.13.711642

**Authors:** Varun B. Dwaraka, Sayf Al-Deen Hassouneh, Kirsten Seale, Dua Sheikh, Jordan Weiter, Joseph Gretzula, Raja Sivamani, Anastasia Georgievskaya, Konstantin Kiselev, Gary M. Fisher, Yilei Cui, Lavinia Propescu, Ryan Smith

## Abstract

Whether distinct visible aging traits, e.g., wrinkling, pigmentation, and inflammation, reflect shared or independent epigenetic programs remains unknown; existing clocks compress aging into a single chronological axis, leaving the phenotype-specific architecture of cutaneous aging uncharacterized. Here, we integrate AI-derived facial phenotypes with skin DNA methylation profiles from 706 individuals to develop EpiVision, a panel of 21 epigenetic predictors spanning structural, pigmentary, inflammatory, and textural aging traits. Predictors reveal shared and trait-specific pathways, including developmental patterning, epithelial remodeling, hormonal signaling, and UV damage responses, and capture environmentally induced acceleration in sun-exposed skin alongside lifestyle and topical treatment–associated variation. These findings establish that visible skin aging comprises molecularly distinct axes with shared regulatory substrates and trait-specific drivers, providing a scalable epigenetic framework for intervention evaluation and aging biology research.

## INTRODUCTION

Human skin aging is a highly visible and clinically relevant phenotype shaped by intrinsic biology and extrinsic exposures. Yet current approaches to measuring skin aging remain fragmented. Imaging-based assessments capture visible structural features, whereas biophysical and molecular assays often require invasive sampling and typically focus on a limited set of endpoints such as elasticity, hydration, collagen content, or barrier function. As a result, existing measures tend to quantify isolated aspects of skin aging rather than providing a comprehensive, multidimensional representation of how skin ages across structural, pigmentary, inflammatory, and textural domains. Moreover, few studies integrate visible phenotypes with molecular readouts or link these measures to longitudinal outcomes.

Recent computer-vision systems trained on large dermatology-labeled image corpora enable standardized, high-throughput extraction of continuous facial skin phenotypes from photographs, offering an opportunity to study skin aging as a multidimensional, quantitative trait at scale^1^. However, these have not been linked to molecular biomarkers, leaving a necessary step for translational research for mechanisms of skin aging.

DNA methylation (DNAm) has emerged as one of the most reproducible molecular readouts of aging across human populations. Early epigenetic clock models demonstrated that DNAm patterns can predict chronological age with high accuracy, establishing DNAm as an integrative biomarker that captures cumulative cellular history across tissues and environmental contexts. However, as DNAm clocks became widely used, it became clear that chronological-age training alone does not guarantee optimal sensitivity to physiology, disease risk, or intervention response. Because first-generation clocks are optimized to minimize error against calendar age, then came the development of second-generation epigenetic clocks trained not on chronological age, but on phenotypes intended to reflect morbidity and mortality risk. DNAm PhenoAge was trained on a composite “phenotypic age” derived from clinical biomarkers and chronological age, and DNAm GrimAge incorporated DNAm surrogates for plasma proteins and smoking exposure to infer biologically meaningful variation that is independent of chronological time, such as heterogeneity in healthspan, tissue-specific decline, or responsiveness to therapeutic intervention. Together, these advances underscored that phenotype-relevant training, rather than chronological age minimization, is necessary to capture biologically meaningful epigenetic variation; a principle that remains largely unapplied in the context of skin aging ^2–5^.

Despite rapid progress in blood-based epigenetic predictors, analogous phenotype-trained DNAm clocks for skin aging remain comparatively underdeveloped. Most widely used skin DNAm clocks remain anchored to chronological time rather than to skin-relevant functional phenotypes. In parallel, the field has begun to develop skin-specific and skin-compatible approaches that lower barriers to sampling and increase sensitivity to cutaneous biology. “MolClock” introduced a skin-focused methylation approach aimed at quantifying skin aging dynamics and treatment effects in a skin-relevant context ^6^. More recently, non-invasive epidermal DNAm profiling from facial tape strips coupled with enzymatic methyl-seq enabled highly accurate chronological age prediction directly from epidermal DNA and demonstrated sensitivity to experimental rejuvenation signals, highlighting the feasibility and translational promise of non-invasive skin epigenomics ^7,8^. Complementing these approaches, newer work has also explored DNAm clocks tied to skin-relevant phenotypes (for example, visual-age progression and wrinkle-grade modeling), underscoring growing interest in phenotype-forward skin aging biomarkers ^9^.

Existing epigenetic clocks vary widely in tissue specificity, training strategy, sample size, and CpG coverage. Early first-generation clocks such as Horvath’s multi-tissue clock (∼8,000 samples; 450K array) ^10^ and the Horvath Skin and Blood clock (∼896 samples) ^11^ were trained primarily on chronological age. Subsequent multi-tissue models, including AltumAge (∼16,000 samples) ^12^ and RetroClock (∼18,000 samples), ^13^ expanded training cohorts and probe coverage but remained largely time-based. Skin-focused approaches such as MolClock (∼508 samples) ^14^, Mitra et al. (epidermal EM-seq; ∼4 million CpGs) ^15^, and commercial second-generation clocks (e.g., Beiersdorf AG) differ substantially in platform depth (450K to >1M CpGs), training size (∼300–800 samples), Cohort demographics (only women) and phenotype focus. Collectively, these models highlight rapid technical progress but also underscore the absence of large-scale, phenotype-trained, imaging-linked skin methylation frameworks grounded explicitly in multidimensional facial aging traits and capable of supporting evidence-based, personalized approaches to long-term skin health.

A key opportunity, and persistent gap, is to connect visible facial aging phenotypes to *molecular* aging signatures in skin using scalable, non-invasive sampling. “Perceived age” from facial images is not merely cosmetic: human-rated perceived age predicts short-term mortality in longitudinal cohorts, indicating that facial appearance can encode health-relevant information beyond chronological age ^16,17^. Consistent with this, blood DNAm has been associated with perceived facial aging, supporting the hypothesis that methylation markers can reflect processes linked to facial aging phenotypes. Advances in deep learning further suggest that facial photographs can support automated estimation of age- and outcome-relevant signals, motivating rigorous efforts to anchor image-derived phenotypes to biological substrates.

A central unresolved question in aging biology is whether the distinct visible features of facial aging (e.g., wrinkling, pigmentation change, inflammation, loss of uniformity) reflect a single shared biological process or independent molecular programs with trait-specific regulation. We address this question directly by integrating AI-derived facial aging phenotypes from HAUT imaging with matched DNA methylation profiles from minimally invasive curette skin samples across 706 individuals. Using this framework, we developed EpiVision, a panel of 21 epigenetic predictors spanning perceived age, structural features, pigmentation, inflammation, and texture, trained on quantitative skin-relevant outcomes rather than chronological time. Alongside these predictors, we performed independent epigenome-wide association analyses for each phenotype to characterize the biological architecture connecting methylation variation to visible aging traits and to quantify the degree of shared versus distinct epigenetic structure across aging axes. Together, these analyses ask not only whether methylation encodes phenotypic information, but which molecular pathways mediate that encoding, and whether different faces of aging are written in the same or different parts of the epigenome.

## RESULTS

### Study collection and design

Facial skin exhibits diverse aging phenotypes that reflect both intrinsic biology and cumulative environmental exposures. To capture DNA methylation signatures underlying these visible features, we developed EpiVision, a methylation-based framework that estimates facial skin aging phenotypes from dermal samples paired with standardized facial imaging. We profiled 744 dermal curette samples collected from the cheekbone region using the Illumina EPICv2 BeadChip, yielding 911,580 CpG sites (97.4% autosomal, 2.6% sex chromosomes) per sample after quality control. In parallel, we analyzed 1,463 standardized facial image submissions and quantified 19 facial aging-related phenotypes using the HAUT artificial intelligence platform. After linkage of methylation and imaging phenotypes, 706 individuals had matched data and were randomly divided into a training set (N=406–564, 80%) and an independent testing set (N=102–142, 20%) with comparable age and Fitzpatrick skin type distributions (**Fig. 1, Supplementary File 1**).

**Figure 1.**
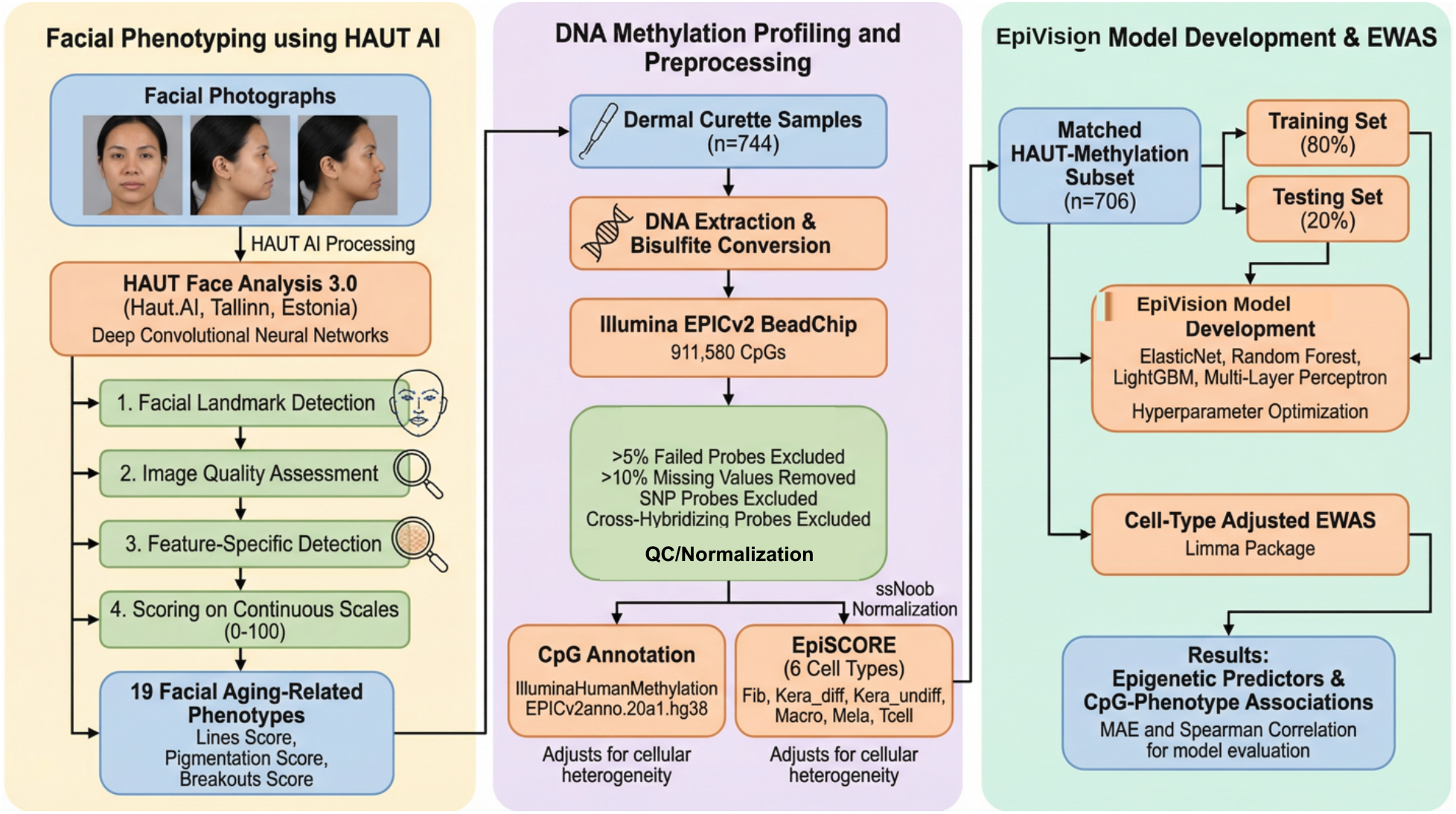
Study design and analytic workflow. Facial photographs were processed with HAUT AI to derive 19 phenotypes (1,463 submissions; n = 1,413 individuals after QC). Facial skin methylation was profiled on the Illumina EPICv2 array (n = 744; 911,580 CpGs after QC). A matched subset (n = 706) was used for two parallel analyses: (i) cell-type-adjusted EWAS of CpG–phenotype associations (EpiSCORE; 6 cell types) and (ii) EpiVision model development using an 80:20 train–test split, evaluated by MAE and Spearman ρ. EpiVision comprises 21 predictors (3 age-related, 18 phenotype-specific). A survey-linked subset (n = 282; 33 exposures) was used for association testing with both unadjusted and age/sex-adjusted models. Sample sizes, algorithm selection, and model details are provided in Supplementary Files 1 and 2.

EpiVision generated a panel of epigenetic predictors that were evaluated in the held-out test set using mean absolute error and Spearman correlation. To examine lifestyle and medical correlates, we linked survey data from 282 participants (33 exposures) and tested exposure– predictor associations using age/sex-adjusted epigenetic age acceleration where applicable. Separately, we performed epigenome-wide association analyses within the matched cohort (N=706) to identify CpG sites associated with specific facial aging features, adjusting for cell-type heterogeneity estimated using EpiSCORE.

### HAUT Imaging provides baseline for skin health and aging

To establish the phenotypic baseline for downstream methylation analyses, we characterized 19 AI-derived facial aging and skin-quality traits across 1,413 individuals from 1,463 imaging submissions using the HAUT platform. The subset with paired methylation data (n = 706) showed highly similar phenotype distributions to the full imaging cohort, indicating representativeness (**Fig. 2A**). Among participants with available demographic data, mean chronological age was 40.3 ± 16.3 years (n = 247), and Fitzpatrick skin types among 481 participants with available data were most commonly Type II (n = 154) and Type III (n = 143) (**Fig. 2B–C**). As expected, structural aging features correlated with chronological age (n = 191–246), with perceived age (r = 0.866) and perceived eye age (r = 0.826) showing the strongest positive associations, while overall uniformness (r = −0.529), lines (r = −0.409), and sagging (r = −0.322) declined with age (**Fig. 2D**). Associations with Fitzpatrick type were also observed (n = 364–468), with irritation (ρ = 0.488) and erythema (HAUT redness score; ρ = 0.470) positively correlated with higher phototypes, and sun spots (ρ = −0.221) and pigmentation score (dyschromia and uneven skin tone; ρ = −0.175) inversely correlated (**Fig. 2E**).

**Figure 2.**
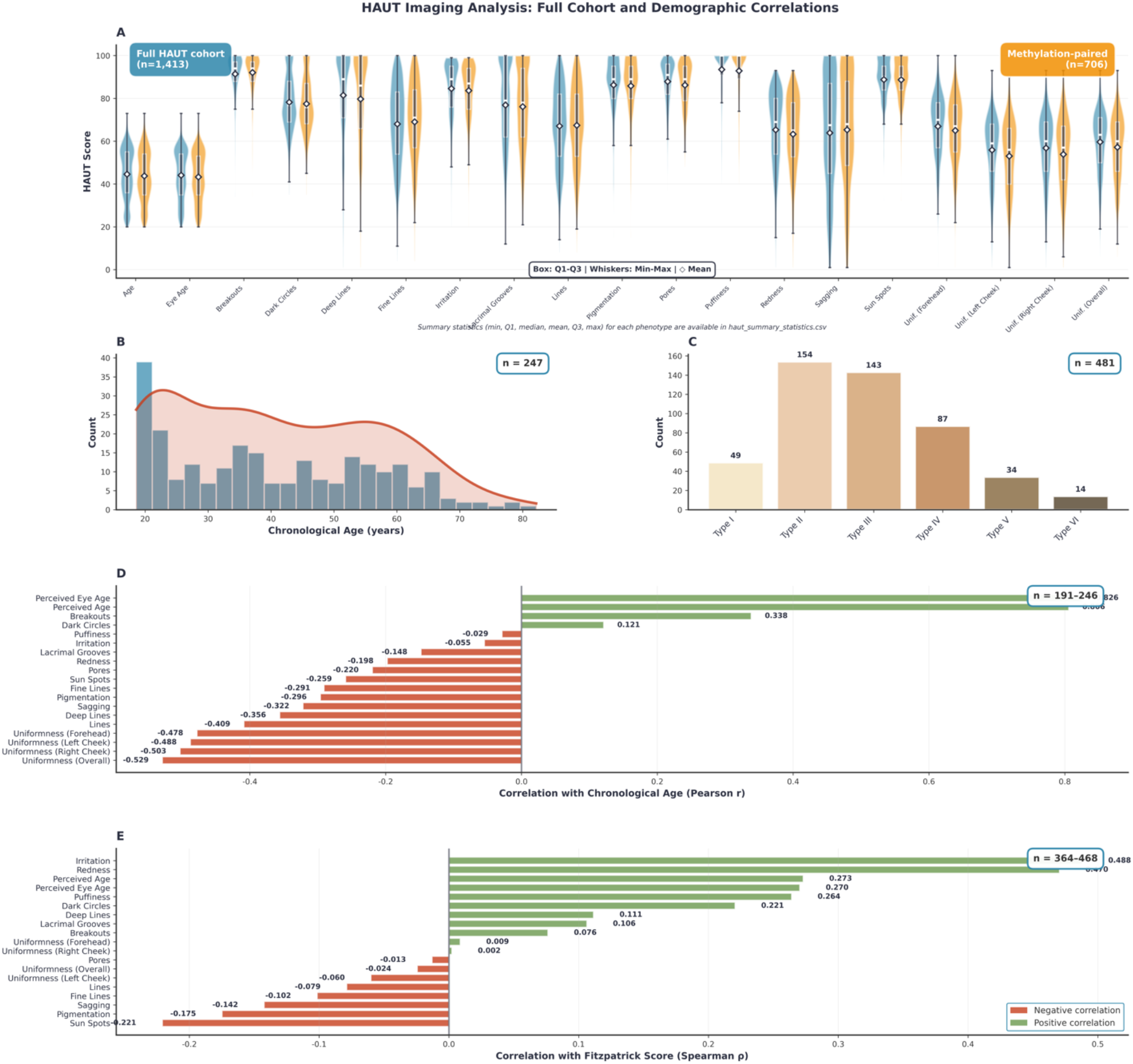
HAUT imaging phenotypes in the full cohort and demographic correlations. (**A**) Distributions of HAUT-derived phenotype scores across the full imaging cohort (blue; n = 1,413) and the subset with paired DNA methylation data (orange; n = 706). Violin plots show score distributions; boxplots indicate Q1–Q3 with median, whiskers denote min–max, and diamonds denote the mean. (**B**) Chronological age distribution for participants with age metadata (n = 247; mean 40.3 ± 16.3 years). (**C**) Fitzpatrick skin type composition for participants with available data (n = 481), with counts shown above bars. (**D**) Pearson correlations between chronological age and HAUT phenotypes (n = 191–246 depending on phenotype). (**E**) Spearman correlations between Fitzpatrick skin type and HAUT phenotypes (n = 364–468). Bar direction and color indicate positive and negative correlations; values are labeled on each bar.

### Skin Cell Deconvolution of methylation data

Cell-type fractions were estimated for 744 samples using EpiSCORE (**Figure S1**). Differentiated keratinocytes represented the largest component (median ≈ 0.32), followed by endothelial cells (median ≈ 0.25) and undifferentiated keratinocytes (median ≈ 0.21). Macrophage and T-cell fractions were present with substantial inter-individual variability, whereas fibroblast and melanocyte fractions were generally low. Overall, the inferred composition, dominated by keratinocytes with a notable endothelial contribution, is consistent with superficial sampling that captures predominantly epidermal cell populations. These cell-type estimates were incorporated as covariates in downstream methylation analyses to account for cellular heterogeneity across samples.

### Development and Performance of Skin Trait Predictors

Skin DNA methylation encodes multidimensional facial aging phenotypes with strong predictive fidelity, demonstrating that the epigenome captures meaningful visible aging information beyond chronological time. To characterize this systematically, we developed epigenetic predictors for 21 skin-related phenotypes using genome-wide DNA methylation profiles from the Illumina EPICv2 platform. Predictors were trained on unadjusted phenotype scores using the full CpG array including sex-linked probes (median 4.8% of selected features per model, range 0.5–33.0%), with sex and age adjustment reserved for downstream association analyses (see Methods). Models for inflammatory traits such as irritation showed the greatest enrichment for sex-linked features, consistent with known sex differences in cutaneous immune regulation. Models were trained in an 80/20 split framework and evaluated in held-out test sets using Spearman correlation and mean absolute error (**Table 1**). Multiple machine learning approaches were assessed, and tree-based ensemble methods consistently outperformed linear models, with LightGBM selected for 17 of 21 predictors (81%) and Random Forest for the remaining four. Age-related predictors showed the strongest performance, including the sun-exposure–informed age model (ρ = 0.836; MAE = 5.30 years), perceived chronological age (ρ = 0.819), and perceived eye age (ρ = 0.765). Several non-age phenotypes also achieved strong correlations, including irritation (ρ = 0.738), uniformness of the left cheek (ρ = 0.709), and redness (ρ = 0.705), demonstrating that methylation profiles capture multidimensional aspects of visible skin biology beyond chronological time.

**Table 1.**
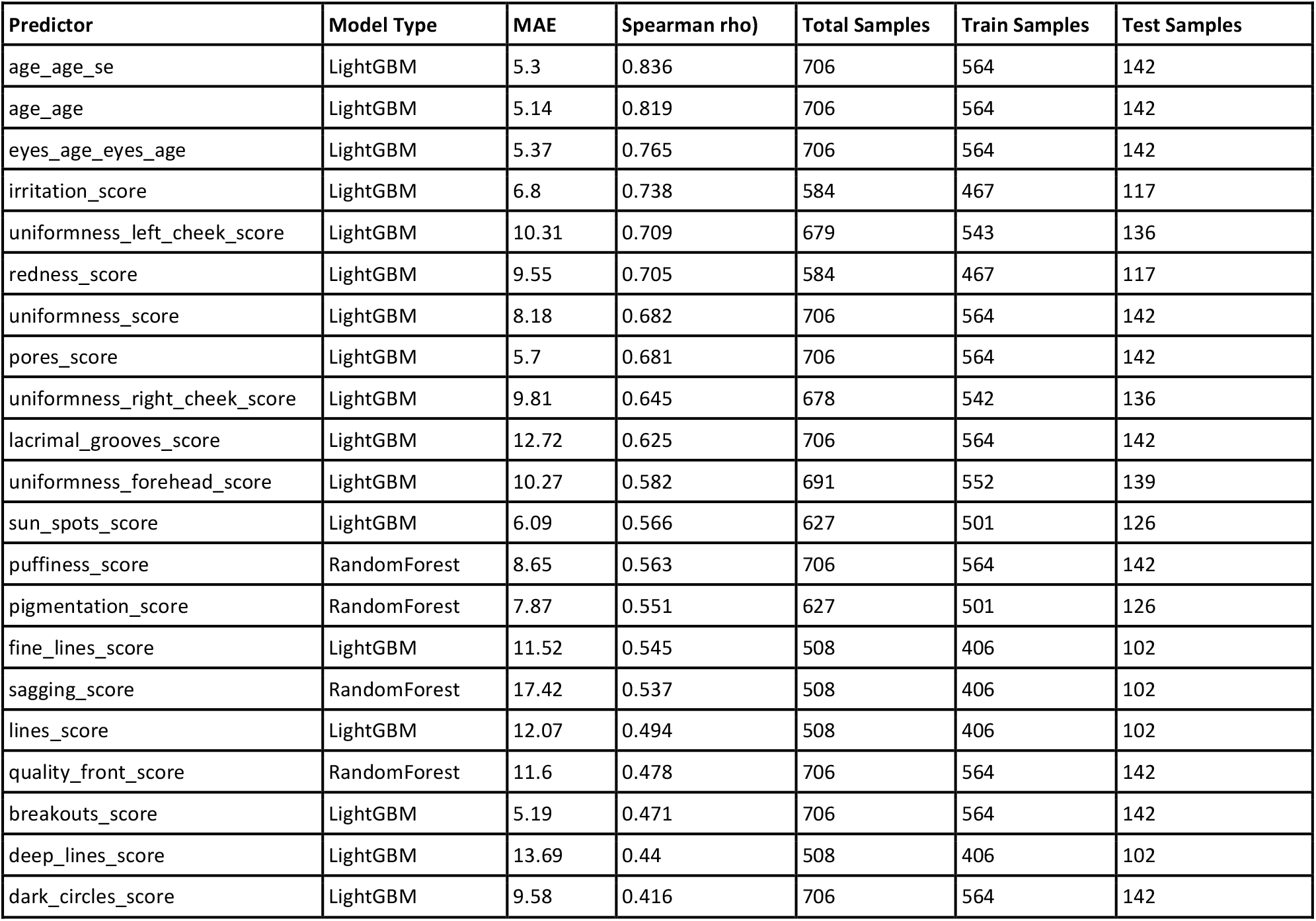
Performance of V2 Models for Skin Trait Prediction. Summary of the final selected model for each of the 21 predictors, sorted by descending Spearman R correlation. MAE (Mean Absolute Error), best Model Type (LightGBM or RandomForest), and sample sizes (total, train, and test) are shown for each predictor.

To evaluate the contribution of expanded probe coverage, we compared EPICv2-specific models with models restricted to CpGs shared between EPICv1 and EPICv2 (**Fig. 3A–D**). EPICv2 models demonstrated superior overall performance, with an average 10.0% increase in Spearman correlation (0.546 vs. 0.601) and 4.6% reduction in MAE (9.83 vs. 9.38) (**Fig. 3A,B**). Applying a ≥5% threshold for meaningful change, EPICv2 models showed meaningful MAE improvement in 45% of targets with no meaningful degradation in any target, and meaningful Spearman improvement in 40% of targets with meaningful degradation in 20% (**Fig. 3C,D**). Improvements were most pronounced for texture-related traits, including lacrimal grooves (ρ = 0.344 to 0.625), overall uniformness, and pores. Although a small subset of predictors (4/20) showed ≥5% decreases in correlation, no models exhibited meaningful increases in MAE. Complete performance metrics for all EPICv2 and V1–V2 overlap models are provided in **Supplementary File 2**.

**Figure 3.**
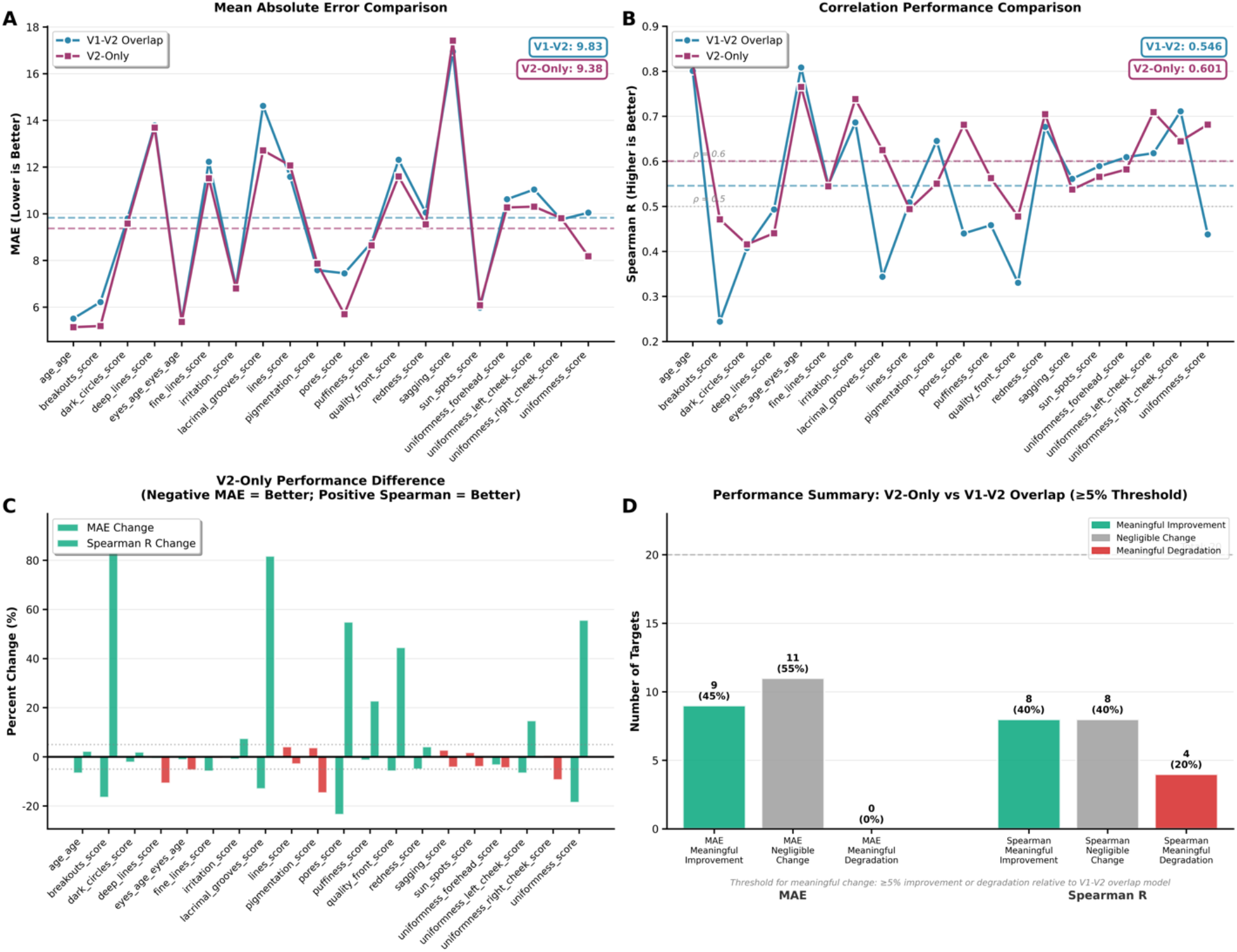
Model performance using EPICv2-only features versus the V1–V2 overlapping CpG set. Performance is shown for 20 targets (chronological age and 19 HAUT phenotypes) trained with identical modeling procedures but different CpG feature spaces. (A) Mean absolute error (MAE; lower is better) for each target. Dashed lines indicate across-target means (V1–V2 overlap: 9.83; V2-only: 9.38). (B) Spearman correlation (ρ; higher is better) for each target. Dashed lines indicate across-target means (V1–V2 overlap: 0.546; V2-only: 0.601). (C) Percent change in performance for V2-only relative to V1–V2 overlap (negative MAE change and positive Spearman change indicate improvement). (D) Summary of meaningful change (≥5% threshold): 9/20 targets (45%) showed meaningful MAE improvement with no meaningful degradation in any target; 8/20 targets (40%) showed meaningful Spearman improvement, with 4/20 (20%) showing meaningful degradation.

### Exposure and Clinical Correlates of EpiVision Predictors

Building on the performance and biological validity of the EpiVision predictors, we examined whether methylation-derived facial aging measures captured expected associations with self-reported lifestyle and clinical factors (**Fig. 4A–B**). Across 282 participants with survey data, exposures were evaluated against 21 methylation-based predictors using age/sex-adjusted regression models to isolate exposure-specific effects from demographic variation captured by the unadjusted predictors, with age-related outcomes analyzed as epigenetic age acceleration (EAA).

**Figure 4.**
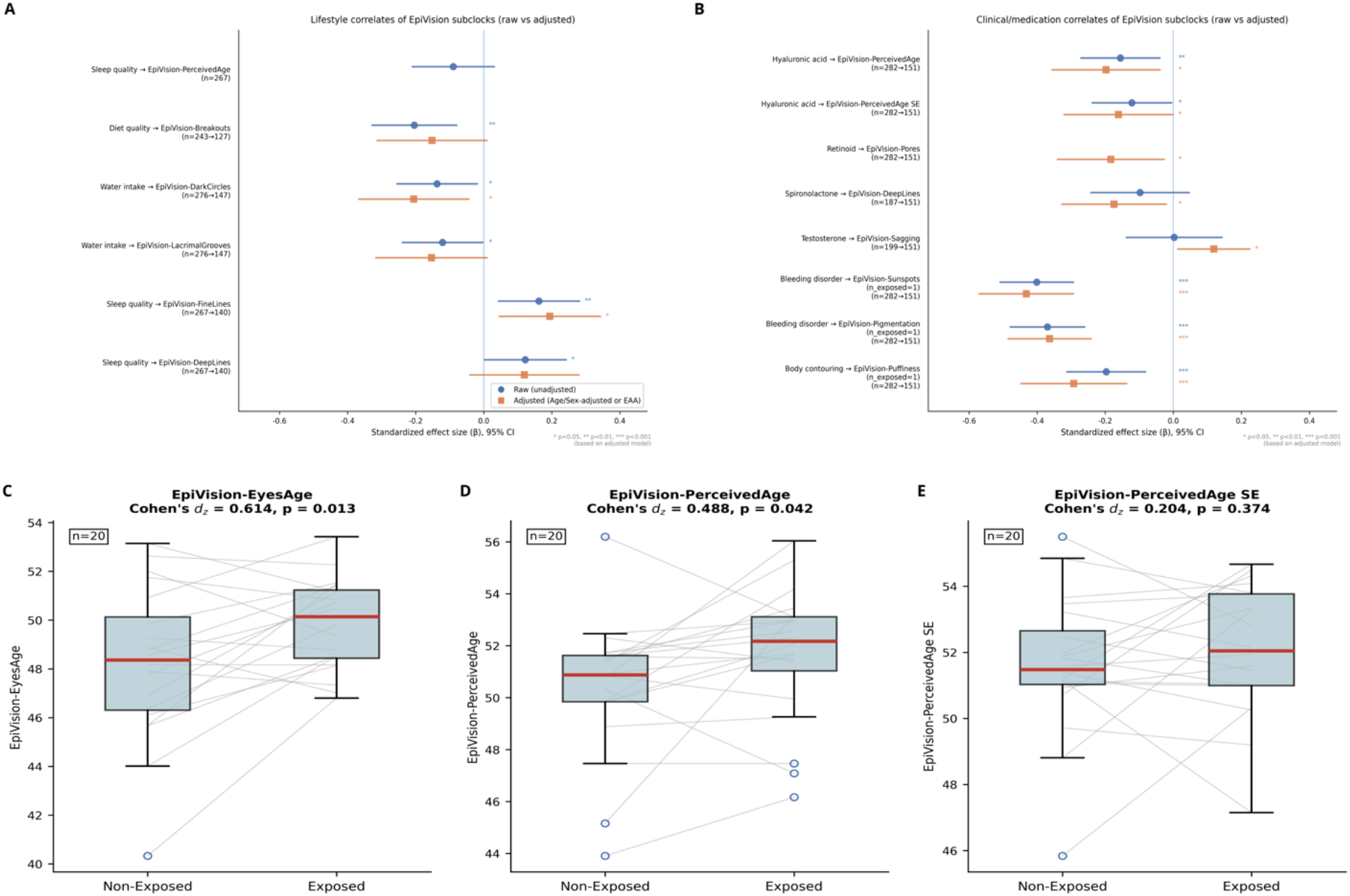
Lifestyle, clinical, and sun-exposure correlates of EpiVision predictors. (A,B) Forest plots showing standardized effect sizes (β) and 95% confidence intervals for associations between self-reported exposures and EpiVision predictors in the survey-linked subset (n = 282). Blue circles denote unadjusted estimates; orange squares denote covariate-adjusted estimates (age/sex-adjusted for phenotype predictors; epigenetic age acceleration for age-related predictors). Sample sizes for each test are shown as n (raw → adjusted), reflecting differences due to covariate availability. (A) Lifestyle correlates. (B) Clinical and medication correlates. Significance asterisks are based on the adjusted model (*P < 0.05, **P < 0.01, ***P < 0.001). (C– E) Paired comparisons of EpiVision age predictions between sun-protected and sun-exposed skin from the same individuals (n = 20). Grey lines connect within-participant measurements. Boxplots show median, interquartile range, and 1.5×IQR whiskers. (C) Eyes Age. (D) Perceived Age. (E) Sun-exposure-informed Perceived Age (SE). Cohen’s d_z and P values from paired t-tests are shown above each panel.

Lifestyle exposures showed phenotype-specific rather than global aging associations, suggesting that behavioral factors influence distinct dimensions of visible skin biology. Greater water intake was associated with reduced dark circles (β = −0.206, p = 0.013), and higher self-reported diet quality was associated with fewer acneiform breakouts (HAUT breakouts score) in the unadjusted model (β = −0.204, p = 0.001), with a consistent direction after age/sex adjustment (β = −0.152, p = 0.065). Sleep quality was significantly associated with fine lines after adjustment (β = 0.194, p = 0.011), with differing effect patterns across line subtypes, indicating that even a single behavioral exposure maps onto structurally distinct aspects of facial aging. Tobacco use demonstrated nominal correlations with pigmentation and line traits in smaller subgroups but did not survive covariate adjustment. The specificity of these associations across phenotypically distinct predictors supports the discriminant validity of the EpiVision framework and highlights the value of decomposing skin aging into trait-level components rather than relying solely on composite aging measures.

Among clinical and medication variables, hyaluronic acid use was the only exposure significantly associated with epigenetic age acceleration, showing slower perceived-age acceleration across both the standard (β = −0.198, p = 0.015) and sun-exposure-informed (β = −0.161, p = 0.048) perceived-age models (**Fig. 4B**). At the phenotype level, retinoid use was associated with reduced pore size appearance (β = −0.184, p = 0.022), spironolactone use with reduced deep line appearance (β = −0.174, p = 0.027), consistent with its known anti-androgenic effects on dermal remodeling, and testosterone use with increased sagging skin (β = 0.119, p = 0.028), an unexpected finding that may reflect residual confounding by age or hormonal context and warrants prospective replication. Notably, unadjusted age analyses revealed numerous significant associations with age-related predictors, including testosterone, hormone therapy, diabetes, and cancer history (all p < 0.005), but these attenuated after EAA adjustment, consistent with confounding by chronological age and reinforcing the necessity of acceleration-based modeling to isolate exposure-specific effects. Many nominal associations attenuated after covariate adjustment, underscoring the influence of age and exposure imbalance, and several large apparent effects were driven by single exposed individuals and are therefore considered exploratory. All association results can be found in **Supplementary File 3**.

To assess the impact of chronic sun exposure on biological aging, we analyzed paired sun-exposed and sun-protected skin samples from 20 participants. The EpiVision perceived-age predictor showed a mean 1.30-year acceleration in sun-exposed tissue relative to sun-protected tissue (paired t-test; Cohen’s d_z = 0.50, p = 0.042; **Fig. 4D**), providing molecular evidence consistent with chronic UV exposure advancing the biological age of skin. The eyes-age predictor showed a directionally consistent and nominally larger acceleration (+1.61 years, p = 0.013; **Fig. 4C**); although trained on periorbital aging phenotypes, the methylation features underlying this predictor are derived from skin broadly and likely capture UV-responsive epigenetic programs shared across cutaneous sites rather than periorbital-specific anatomy.

Beyond epigenetic acceleration, sun exposure broadly altered visible skin phenotypes (**Supplementary File 4**). Nine of 18 non-age traits (50%) differed significantly between sites, with the largest effects observed for erythema, sagging skin, pigmentary change, and lines, all demonstrating large effect sizes (|d| > 0.8). Acneiform breakouts were more prevalent in sun-exposed skin, consistent with UV-induced disruption of barrier integrity, increased sebaceous activity, and follicular occlusion in photodamaged tissue. The convergence of structural, pigmentary, and inflammatory changes across sites indicates that chronic UV exposure perturbs multiple biological axes simultaneously rather than isolated features. Together, these paired-sample analyses demonstrate that chronic sun exposure accelerates both molecular epigenetic aging and a broad spectrum of visible skin phenotypes, and establish the EpiVision perceived-age predictor as a biologically grounded, skin-appropriate molecular readout of environmentally induced aging.

### Identification of differentially methylated positions associated with facial skin features

Having established that skin methylation encodes phenotype-level aging information, we next asked what biological processes underlie this connection and whether distinct visible aging traits share a common epigenetic substrate or arise from independent molecular programs. To address this, we conducted a parallel, independently designed epigenome-wide association study (EWAS) for each of the 19 HAUT phenotypes. Critically, this analysis was structured to adjust for confounders that predictive models absorb implicitly but do not decompose: technical batch effects (Illumina BeadChip), inter-individual variation in skin cell-type composition (EpiSCORE; six fractions), and biological sex. This covariate-adjusted design isolates trait-specific epigenetic signals from artifacts of cellular heterogeneity and demographic imbalance. The number of significant DMPs varied substantially across traits, ranging from 3,050 for lacrimal grooves (0.33% of tested sites) to 367,506 for acneiform breakouts (40.3%), reflecting marked differences in epigenetic architecture across phenotypes.

To characterize the shared epigenetic architecture underlying these phenotypes, we performed a cross-trait pleiotropy analysis across all 19 EWAS traits (Supplementary File 5; two traits from the original 21-trait panel, perceived age standard error and quality front, were excluded from the EWAS owing to insufficient phenotypic variance after quality filtering). Of 586,859 unique CpG sites reaching significance (FDR < 0.05) in at least one trait, 43.5% were trait-specific, 48.4% were significant in two to four traits, and 7.8% in five to nine traits. Only 1,483 CpGs (0.25%) were significant in ten or more traits, a distribution that departs dramatically from the expectation under independence (χ^2^ = 2,118,347, df = 3, P < 10^−300^). This small set of highly pleiotropic loci, significant across more than half the phenotype panel, represents a shared epigenetic substrate of facial aging. Of these, 55.0% mapped to intergenic regions, suggesting that long-range regulatory elements play a substantial role in coordinating multi-trait aging biology. Pairwise Jaccard similarity of significant CpG sets revealed biologically coherent clustering: perceived age and perceived eye age shared the greatest overlap (J = 0.78), the three uniformness sub-regions formed a tight cluster (J = 0.41–0.62), pigmentation and sun spots (solar hyperpigmentation) grouped together (J = 0.47), lines and sagging co-clustered (J = 0.37), and irritation and erythema formed a distinct inflammatory module (J = 0.31). Nearly all highly pleiotropic CpGs (99.8%) showed mixed directionality, with hypermethylation in some traits and hypomethylation in others, rather than uniform effects. Because all models adjusted for six EpiSCORE cell-type fractions and technical batch, this mixed directionality is unlikely to reflect compositional confounding and instead suggests that pleiotropic loci participate in genuinely distinct, trait-specific regulatory programs.

These multi-trait loci represent candidate targets for future functional and intervention studies: modulating a pleiotropic locus could, in principle, exert coordinated effects across structural, pigmentary, and inflammatory aging axes simultaneously. The most broadly associated CpG was cg16867657 at the ELOVL2 locus, significant in 15 of 19 traits spanning deep lines, fine lines, perceived age, pigmentary change, erythema, sagging, and all four uniformness measures, among others. Other highly pleiotropic loci included cg08870112 (RANBP17; 14 traits), cg07553761 (TRIM59; 14 traits), and cg23571812 (FOXG1; 13 traits), all established epigenetic aging biomarkers. Recurrent gene annotations among the 1,483 highly pleiotropic CpGs included ZIC1 (13 CpGs), PRRT1 (11 CpGs), ZNF154 (7 CpGs), and FOXG1 (6 CpGs), implicating developmental transcription factors and zinc-finger proteins in the shared architecture of skin aging. The direction of effect also differed systematically by phenotype category: structural aging features such as lines and sagging showed predominantly hypermethylated DMPs, consistent with progressive locus-specific gain of methylation with aging, while inflammatory traits including irritation and acneiform breakouts showed predominantly hypomethylated positions, suggesting loss of epigenetic regulation at immune-relevant loci. Full DMP results for all 19 traits are provided in **Supplementary File 5**.

### Perceived age and perceived eye age

For perceived age (ρ = 0.82), enrichment was dominated by neuronal development and morphogenesis terms including axonogenesis (FDR < 10^−22^), alongside epithelial–mesenchymal transition (EMT; FDR = 3.43×10^−4^), apical junction, NOTCH signaling, and estrogen response programs; this implicates coordinated tissue-architectural remodeling rather than isolated molecular damage. Many enriched neuronal guidance genes (SLIT2, ROBO1/2, SEMA3A/F, NCAM1) also have established roles in cutaneous innervation and epithelial organization, underscoring shared developmental programs between neural and skin tissues. Perceived eye age (ρ = 0.765) showed a largely overlapping architecture, with developmental patterning and axonogenesis (FDR < 10^−24^), EMT, and estrogen response as top enriched terms, with additional myogenesis enrichment consistent with periorbital muscle–skin structural interactions. The substantial overlap between these two aging phenotypes, corroborated by their exceptionally high Jaccard similarity (J = 0.78), indicates that shared developmental and structural epigenetic programs underlie aging across anatomically distinct facial regions.

#### Additional traits (ρ > 0.7)

Irritation (ρ = 0.74) highlighted innate immune activation, cytokine production, and neurogenesis regulation pathways, pointing to epigenetic encoding of inflammatory skin states. Redness (ρ = 0.70) showed enrichment for neurogenesis regulation and vascular processes including angiogenesis and vascular permeability, reflecting the neurovascular biology underlying cutaneous erythema. Uniformness (left cheek; ρ = 0.71) implicated epithelial organization and keratinocyte turnover programs. Traits with weaker epigenetic–phenotypic correlations (ρ < 0.70), including uniformness overall (ρ = 0.68), pores (ρ = 0.68), lacrimal grooves (tear trough areas; ρ = 0.63), and pigmentation (ρ = 0.55), showed analogous but attenuated enrichment themes involving developmental signaling and epithelial architecture. For pigmentation, despite the more modest predictive correlation, pathway enrichment revealed UV-response and melanocyte morphogenesis programs consistent with photoaging-driven dysregulation of pigment biology, including both hyper- and hypopigmentation.

Across all traits, developmental signaling, epithelial junction biology, estrogen response, and UV-associated programs recurred as shared substrates of facial aging, while trait-specific signals, such as UV response in pigmentary change, metabolic stress in irritation, and vascular programs in erythema, indicate layered biological specialization atop this common aging architecture. CpG loci and pathways enriched across multiple phenotypes, including the developmental and epithelial-junction programs shared by perceived age, uniformness, and erythema, may represent particularly actionable intervention targets, as modulating them would be expected to influence multiple aging dimensions simultaneously. Complete pathway enrichment results across GO, KEGG, and Hallmark databases (28,583 enriched terms across 19 traits) are provided in **Supplementary File 6**.

## DISCUSSION

Visible human facial aging is not a single biological process. The findings presented here demonstrate that the skin epigenome encodes distinct molecular programs underlying structural aging, pigmentary change, inflammation, and textural change, programs that are partially shared across traits but that also carry trait-specific biological signatures. By integrating AI-derived facial phenotypes with matched skin DNA methylation profiles across 706 individuals, EpiVision establishes that methylation variation in superficial skin samples captures multidimensional aging information with biological coherence, environmental sensitivity, and responsiveness to clinical exposures. Three primary advances emerge: skin methylation accurately predicts specific facial aging dimensions well beyond what chronological age alone explains; trait-specific epigenome-wide analyses reveal biologically coherent and partially independent molecular architectures underlying each visible aging axis; and EpiVision predictors capture expected relationships with UV exposure, lifestyle factors, and dermatologic interventions, validating both their biological grounding and practical utility.

A particularly compelling observation concerns the relationship between EpiVision’s trait architecture and clinically recognized photoaging subtypes. Sachs et al. characterized two divergent forms of UV-induced skin aging: hypertrophic photoaging, marked by coarse wrinkling, skin thickening, and sallowness, and atrophic photoaging, characterized by fine wrinkling, erythema, telangiectasia, and elevated keratinocyte cancer risk ^18^. Subsequent histological work confirmed distinct structural features underlying these phenotypes, including differential solar elastosis and collagen VII distribution ^19^. EpiVision’s EWAS enrichment signatures map onto this clinical axis with notable specificity. The erythema and irritation predictors showed neurogenesis regulation among their most significantly enriched processes, alongside innate immune activation, vascular processes, and angiogenesis. This profile mirrors the neurogenic and vascular inflammatory biology described as hallmarks of atrophic photoaging. Conversely, the lines, sagging, and deep lines predictors were enriched for extracellular matrix organization, collagen fibril organization, and fibroblast migration, reflecting the structural ECM remodeling and elastotic thickening that define the hypertrophic phenotype. Although the current cohort lacks formal AP/HP clinical classification, the convergence of these orthogonal enrichment signatures suggests that EpiVision’s trait architecture may already capture the molecular underpinnings of this clinical dichotomy. Prospective integration of AP/HP classification with EpiVision outputs would provide a framework for testing whether distinct epigenetic signatures predict photoaging subtype or modulate responsiveness to subtype-specific interventions.

A key insight from this work is that skin DNA methylation encodes biologically meaningful information about visible aging phenotypes beyond what is predicted by chronological age. This connection has not previously been demonstrated at this phenotypic resolution. Age-related predictors achieved robust test-set correlations (sun-exposure–informed model, ρ = 0.836; perceived age, ρ = 0.819; eye age, ρ = 0.765), while non-age phenotypes including irritation, erythema, and regional uniformness exceeded ρ = 0.70, indicating that methylation captures multidimensional cutaneous biology beyond chronological time.

The parallel EWAS analyses, performed independently of model training and adjusted for batch effects, cell-type composition, and sex, provide biological grounding for the predictive performance. For phenotypes with strong epigenetic coupling, enrichment analyses converged on developmental and structural pathways. Perceived age, for instance, was enriched for neuronal development alongside epithelial–mesenchymal transition, apical junction, and estrogen response programs, implicating coordinated tissue-architectural remodeling rather than passive damage accumulation. Several enriched neuronal guidance genes, including SLIT2, ROBO1/2, SEMA3A/F, and NCAM1, also play established roles in cutaneous innervation and epithelial organization, underscoring shared developmental programs between neural and skin tissues. Trait-specific signals aligned with known dermatologic drivers, including UV-response pathways for pigmentation and innate immune activation for irritation. Cross-trait pleiotropy analysis identified shared “core” aging loci: cg16867657 (ELOVL2) was significant in 15 of 19 traits, consistent with the established role of ELOVL2 methylation as a pan-tissue aging marker ^20^. The direction of effect also differed systematically. Structural features showed predominantly hypermethylated DMPs consistent with progressive locus-specific methylation gain, while inflammatory traits showed predominantly hypomethylated positions suggesting loss of epigenetic regulation at immune-relevant loci.

We validated EpiVision’s biological sensitivity through paired sun-exposure analysis and survey-linked lifestyle associations. In 20 participants with matched sun-exposed and non–sun-exposed samples, sun-exposed skin showed accelerated perceived age (+1.30 years, p = 0.042) alongside deterioration in 9 of 18 phenotypic traits. In the lifestyle analyses, exposures showed phenotype-specific rather than global aging associations: water intake was linked to reduced dark circles, sleep quality to fine line measures, and self reported topical hyaluronic acid use was the only exposure significantly associated with epigenetic age acceleration across both perceived-age models. This pattern, where lifestyle factors influence individual trait predictors rather than shifting a composite clock, reinforces the value of decomposing skin aging into trait-level components and positions EpiVision as a more informative endpoint than single-score approaches. Notably, numerous raw associations with age-related predictors (testosterone, HRT, diabetes, cancer history) attenuated after EAA adjustment, reinforcing the necessity of acceleration-based modeling to isolate exposure-specific effects from chronological age confounding.

EpiVision extends the phenotype-forward direction introduced by Bienkowska et al., who developed VisAgeX, a skin-specific second-generation epigenetic clock predicting visual age progression from epidermal methylation profiles of female volunteers ^21^. While VisAgeX targeted wrinkle grade and visual facial age using 450K-array data from a single-sex cohort, EpiVision broadens both the phenotypic scope (21 predictors spanning structural, pigmentary, inflammatory, and textural axes) and population diversity (both sexes, multiple Fitzpatrick types) while leveraging EPICv2 coverage for enhanced regulatory CpG representation. EpiVision also complements earlier skin-specific chronological-age clocks ^22^, senotherapeutic validation platforms ^23^, and emerging non-invasive epidermal methylomics approaches, including tape-strip whole-genome profiling ^24^ and sequencing-based epigenetic age predictors optimized for non-invasively collected epidermis ^25^. Together, these advances position skin methylation as an increasingly tractable molecular layer for both aging research and intervention evaluation.

Several limitations warrant consideration. The cross-sectional design precludes causal inference and assessment of within-person trajectories; longitudinal tracking will be essential to quantify intervention responsiveness and determine whether EpiVision captures true biological change over time. The cohort’s ancestry and Fitzpatrick diversity is limited (Types V–VI comprised 6.5% of the methylation cohort), and expansion to more diverse populations is needed to establish generalizability. Integration with transcriptomic and proteomic layers would connect methylation shifts to functional outputs and strengthen mechanistic interpretation. Continued refinement of deconvolution and sampling protocols will help reduce residual confounding by cellular heterogeneity. Biophysical measures such as transepidermal water loss (TEWL) and skin hydration were not captured in the current study and would complement methylation-derived readouts in future work. Future work should establish whether EpiVision predictors capture true biological trajectories across diverse populations and serve as scalable endpoints for mechanistically informed, data-driven skin-aging models. More broadly, the demonstration that skin methylation encodes biologically distinct aging axes, rather than a single chronological signal, opens a path toward understanding facial aging as a decomposable biological phenotype, with implications for how epigenetic tools are designed, validated, and applied across aging research, clinical dermatology, and regenerative dermatologic practice.

## METHODS

### Study Population and Sample Collection

Facial skin samples were collected from participants enrolled in a skin aging cohort, including the TruDiagnostic cohort, Integrative Skin Research (led by Dr. Raja Sivamani), and Boynton Beach Dermatology / Beauty Within (led by Dr. Joseph C. Gretzula), using sterile dermal curettes (Integra Miltex, Integra LifeSciences), an instrument designation reflecting the tool rather than the tissue depth, to obtain predominantly epidermal skin cells, as confirmed by EpiSCORE deconvolution. Skin samples were collected from the cheekbone region and subsequently transferred to collection vials containing DNA/RNA Shield stabilization buffer (Zymo Research, Irvine, CA) and shipped to the laboratory for processing. DNA was extracted from collected skin samples using standard protocols and quantified prior to bisulfite conversion.

Participants completed questionnaires, facial images, and provided informed consent prior to participation. The questionnaires captured demographic information as well as relevant skin history, health history, and potential environmental exposures. All study procedures were approved by the relevant institutional review boards. The final analytic sample comprised 508 to 706 individuals per phenotype depending on data availability (see Results for phenotype-specific sample sizes).

### Facial Phenotyping Using HAUT Artificial Intelligence Platform

#### Image Acquisition

Participants submitted standardized facial photographs (selfies) using their personal mobile devices. Images were captured in three standard orientations: frontal view, left profile, and right profile. Participants were instructed to photograph themselves in well-lit environments without makeup, with hair pulled back to expose the full face, and with neutral facial expressions. Multiple submissions from the same participant were allowed over time to accommodate technical failures or quality issues.

#### Automated Phenotype Extraction

All facial images were processed using HAUT Face Analysis 3.0 software (Haut.AI, Tallinn, Estonia), an artificial intelligence-powered computer vision platform designed for automated, objective assessment of skin aging and appearance characteristics from digital photographs. The HAUT platform employs deep convolutional neural networks trained on large-scale annotated datasets of facial images labeled by expert dermatologists for a comprehensive array of skin features.

For each submitted image, the HAUT pipeline performs the following automated steps: (1) facial landmark detection to identify anatomically defined regions of interest (forehead, periorbital area, cheeks, perioral region, nose); (2) image quality assessment across multiple technical dimensions including resolution, blur, exposure, illumination uniformity, proper facial positioning, absence of occlusions (eyeglasses, hair, hands), and sufficient visible skin area; (3) feature-specific detection using specialized deep learning models for each phenotype (e.g., wrinkle detection models identify line segments and quantify depth, length, and spatial distribution; pigmentation models segment hyperpigmented lesions and compute density and color metrics; inflammatory lesion models identify and count pimples, papules, and pustules); and (4) scoring on continuous scales (0-100 for most traits, with higher scores generally indicating better skin quality or less severe manifestation; or estimated age in years for perceived aging phenotypes).

The HAUT platform generates the following phenotypic outputs that were used in the current epigenome-wide association studies: (1) aging-related phenotypes including chronological age (self-reported age in years) and perceived eye age (AI-estimated apparent age of the periorbital region based on comparison to age-labeled reference datasets); (2) structural aging phenotypes including general lines score (overall facial wrinkle burden), fine lines score (shallow wrinkles <0.5mm depth), deep lines score (pronounced wrinkles >1mm depth), sagging score (gravitational descent of facial soft tissues and jowl formation), and lacrimal grooves score (infraorbital hollowing and tear trough deformity); (3) pigmentation phenotypes including overall pigmentation score (dyschromia and uneven skin tone) and sun spots score (discrete solar lentigines and hyperpigmented macules); (4) inflammatory and skin quality phenotypes including breakouts score (acneiform lesions, pustules, papules), irritation score (erythema, scaling, roughness), and redness score (vascular prominence and telangiectasia); (5) textural phenotypes including pores score (visible pore size and density) and puffiness score (periorbital edema); (6) additional localized features including dark circles score (infraorbital hyperpigmentation and shadowing); and (7) skin tone uniformness assessed globally (overall uniformness score) and regionally (forehead, left cheek, right cheek uniformness scores measuring color heterogeneity and blotchiness within each facial zone).

### Quality Control and Best-Row Selection

Embedded quality control algorithms within the HAUT platform assigned composite quality scores to each image based on the aforementioned technical criteria. For participants with multiple image submissions, we selected a single “best-row” image per individual defined as the submission with the highest composite quality score, thereby maximizing measurement reliability while avoiding pseudoreplication from repeated measures. Images failing to meet minimum quality thresholds were excluded from analysis. This quality control process resulted in the retention of 1,413 to 1,463 individuals with usable HAUT phenotype data depending on the specific trait (see Results for details).

### DNA Methylation Profiling and Preprocessing

#### Array-Based Methylation Measurement

Genomic DNA extracted from skin curette samples was bisulfite-converted using the EZ DNA Methylation Kit (Zymo Research, Irvine, CA, USA) according to the manufacturer’s protocol. Bisulfite-converted DNA was hybridized to the Illumina Infinium MethylationEPIC v2 BeadChip (Illumina, San Diego, CA, USA), which interrogates DNA methylation at over 935,000 CpG sites across the human genome with enhanced coverage of regulatory regions including gene promoters, CpG islands, enhancers, and CTCF binding sites. Arrays were processed following the Illumina standard protocol, with samples randomly distributed across multiple BeadChips (recorded as the “Beadchip” batch variable) to mitigate batch effects. Fluorescence intensity data (IDAT files) were generated using the Illumina iScan System.

#### Quality Control

Raw IDAT files were imported into R version 4.4.0 (R Core Team, 2024) and processed using the minfi Bioconductor package (Aryee et al., 2014). Quality control was performed using detection p-value filtering with the following criteria: (1) samples with mean detection p-value >0.15 across all probes were excluded; (2) probes with mean detection p-value >0.10 across all samples were removed (n = 19,078 probes excluded). Sex chromosome probes were retained in the methylation feature matrix, as X- and Y-linked CpG methylation captures biologically relevant sex-differential variation in skin biology (15,722 chrX probes, 46 chrY probes retained).

#### Data Normalization

Normalization was performed using the ssNoob (single-sample normal-exponential out-of-band) method implemented in the preprocessNoob function with dyeMethod = “single” (Triche et al., 2013), which models background correction and dye-bias normalization in a single unified framework using out-of-band probe intensities. The ssNoob method performs within-array normalization, making it suitable for datasets where samples are processed in multiple batches and providing robustness against batch effects. For epigenetic clock prediction, missing values at required clock CpG sites were imputed using reference-based imputation implemented in the SeSAMe package (Zhou et al., 2018).

#### Beta extraction

Following quality control and normalization, 911,580 CpG sites remained for downstream epigenome-wide association analyses, including both autosomal and sex chromosome probes. Methylation levels were represented as beta values (β = M / [M + U + 100], where M and U represent methylated and unmethylated signal intensities, respectively), ranging from 0 (completely unmethylated) to 1 (completely methylated). Genomic coordinates are reported relative to the hg38 human genome reference build (EPICv2 custom array annotation version 20a1).

#### CpG Annotation

CpG sites were annotated using the IlluminaHumanMethylationEPICv2anno.20a1.hg38 R package, which provides gene annotations (mapping to UCSC gene symbols via proximity to transcription start sites), genomic locations (chromosome and position), and regulatory context (CpG island, shore, shelf, or open sea classification). Intergenic CpG sites (those not mapping to annotated gene bodies or promoters within 1.5 kb of transcription start sites) were labeled as “intergenic” throughout the Results.

#### Cell Type Deconvolution for Adjustment of Cellular Heterogeneity

Skin is a heterogeneous tissue comprising multiple cell types with distinct DNA methylation profiles, including keratinocytes, fibroblasts, melanocytes, endothelial cells, and resident immune cells. To distinguish true within-cell methylation changes from artifacts arising due to inter-individual differences in skin cell-type composition, we estimated cell-type proportions from the DNA methylation data using a reference-based deconvolution approach.

Cell type fractions were estimated using the EpiSCORE algorithm (Teschendorff et al., 2017) with the skin-specific methylation reference matrix mrefSkin.m, as implemented in the EpiSCORE R package version 1.0. Although EpiSCORE was originally developed for blood, we applied the skin-specific reference matrix mrefSkin.m because our samples are derived from skin tissue and the skin-specific reference captures the relevant cell-type heterogeneity in our dataset. The mrefSkin.m reference was derived from purified cell populations isolated from human skin tissue and includes methylation signatures for six cell types: fibroblasts (Fib, dermal cells responsible for extracellular matrix production), differentiated keratinocytes (Kera_diff, terminally differentiated epidermal barrier cells), undifferentiated keratinocytes (Kera_undiff, progenitor keratinocytes in the basal layer), macrophages (Macro, tissue-resident immune cells), melanocytes (Mela, pigment-producing cells), and T cells (Tcell, adaptive immune cells).

The EpiSCORE deconvolution procedure consists of two steps. First, CpG-level methylation data (beta values) are aggregated to gene-level methylation summaries by averaging CpG sites within 200 base pairs of annotated transcription start sites (TSS) using the constAvBetaTSS function with the “850k” array type specification (compatible with EPICv2 annotation). This gene-level aggregation reduces technical noise and increases the robustness of deconvolution by focusing on regulatory regions most likely to differ across cell types. Second, the gene-level methylation matrix is input to the weighted robust partial correlation (wRPC) function, which estimates cell type proportions by comparing the sample methylation profile to the reference cell type signatures while down-weighting unreliable genes (useW = TRUE, weight threshold wth = 0.4). This approach is robust to batch effects and outliers. The resulting cell type fraction estimates (proportion of each cell type per sample, summing to 1 across the six cell types) were merged with phenotypic data and included as covariates in all epigenome-wide association models.

### Development of skin predictors

We developed a two-stage framework to identify robust and biologically interpretable DNA methylation features and to train and validate predictive models for multiple biomarker targets using complementary machine learning architectures with systematic hyperparameter optimization. The workflow was designed to capture complex, potentially non-linear and non-monotonic relationships between CpG methylation and disease-relevant phenotypes while preserving directional interpretability for selected features.

To quantify feature importance with a conservative selection threshold, we implemented a hybrid feature-importance procedure that integrates a noise-based baseline, repeated ensemble-derived importance estimation, and post hoc directionality inference. A synthetic Gaussian noise feature was generated using the empirical distribution of the input data and appended to the feature matrix to establish a minimum importance “floor.” During model fitting, this noise feature was treated identically to biological features, enabling prioritization of CpG sites whose importance exceeded that of the noise feature and reducing retention of spurious predictors. Feature importance was estimated using random forest regression under repeated instantiation to reduce variance due to stochastic training effects and mitigate overfitting during feature selection. Specifically, the random forest model was refitted 100 times using independent random seeds and aggregated importance values were used to identify features that were consistently informative across runs. Because tree-based importance measures do not preserve the direction of feature effects, directional relationships for retained features were inferred using multiple linear regression fit to the selected feature set, and the sign of each regression coefficient was used to assign positive or negative directionality relative to the modeled target. This combined approach supports global feature evaluation rather than pairwise screening, captures non-linear and non-monotonic relationships through ensemble modeling, and improves biological interpretability by providing signed effect estimates that clarify how each feature contributes to age estimation, disease prediction, or biomarker quantification.

To identify robust predictive signatures in DNA methylation data, we implemented a comprehensive model development and validation framework that evaluates multiple regression and classification algorithms under rigorous hyperparameter optimization. For each target, four model families were trained and compared: ElasticNet, random forest regression, LightGBM gradient boosting, and a multi-layer perceptron. This set was chosen to span regularized linear models, bagged tree ensembles, gradient-boosted trees, and neural networks, reflecting prior observations that optimal modeling architectures can vary across biomarkers as a function of underlying data dispersion and distributional characteristics.

All models underwent systematic hyperparameter tuning using the Optuna framework to efficiently navigate complex search spaces while limiting overfitting. ElasticNet models were optimized across the regularization strength and L1 ratio to identify sparse yet stable methylation signatures. Random forest regressors were tuned across the number of estimators, maximum depth, minimum samples required to split, and feature selection settings. LightGBM models were tuned across learning rate, number of leaves, tree depth, feature and bagging fractions, and regularization parameters. The neural network architecture was optimized over the number of hidden layers, nodes per layer, activation function, learning rate, and batch size.

Model performance was evaluated using a standard hold-out validation strategy with a random 80/20 train–test split. Models were trained on unadjusted HAUT phenotype scores to preserve the full range of biologically meaningful variation, including sex-dependent and age-dependent differences in skin aging; biological sex and chronological age were instead included as covariates in all downstream association and EWAS analyses to isolate exposure- and trait-specific effects. Predictive performance was quantified using mean absolute error to assess absolute accuracy and Spearman correlation to assess rank preservation between predicted and observed values. Final model selection was based on minimizing a composite objective in which absolute error was penalized based on Spearman’s rho, thereby balancing accuracy with correlation strength and prioritizing models that were both accurate and rank-consistent for the outcome of interest.

### Epigenome-Wide Association Studies: Statistical Framework

#### Data Preparation and Transformation

For each of the 19 HAUT-derived phenotypes, samples with missing phenotype values were excluded from the analysis. The remaining samples were subset from the full methylation matrix, and beta values were logit-transformed to M-values using the formula M = log_2_(β / [1 - β]) to better satisfy the assumptions of linear regression, as M-values are more homoscedastic (constant variance) across the methylation range compared to beta values (Du et al., 2010). CpG sites with any infinite or missing M-values (which can occur if beta values are exactly 0 or 1 due to technical artifacts) were excluded from the analysis.

#### Differential Methylation Analysis

Differential methylation analysis was performed using the limma package version 3.62.1 (Ritchie et al., 2015), which implements linear models for each CpG site with empirical Bayes moderation of standard errors to improve statistical power, particularly for small to moderate sample sizes. For each phenotype, we fit the following linear model at each CpG site:

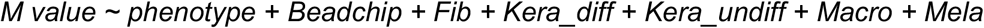

where M represents the M-value (logit-transformed methylation: log_2_[β/(1-β)]) at a given CpG site; phenotype is the continuous HAUT-derived score (0–100 scale), representing the primary independent variable of interest; Beadchip is a categorical variable (factor) representing the Illumina BeadChip identifier on which the sample was processed, included to adjust for technical batch effects; and EC, Fib, Kera_diff, Kera_undiff, Macro, and Mela are continuous numeric variables representing the estimated cell type fractions (proportions) for endothelial cells, fibroblasts, differentiated keratinocytes, undifferentiated keratinocytes, macrophages, and melanocytes, respectively, included to adjust for confounding due to inter-individual variation in cellular composition. All models used treatment (dummy) coding for categorical variables, as specified by options(contrasts = c(“contr.treatment”, “contr.treatment”)) in R. Note: Tcell (T cells) was initially included but was non-estimable across all phenotypes due to perfect collinearity with other cell type fractions and was automatically dropped by the model fitting algorithm.

The linear model was fit using the `lmFit` function, which efficiently estimates regression coefficients across all CpG sites simultaneously via matrix operations. Empirical Bayes shrinkage of residual variances was then applied using the `eBayes` function, which borrows information across CpG sites to stabilize variance estimates and compute moderated t-statistics. Differential methylation statistics for the phenotype coefficient (logFC, t-statistic, unadjusted P-value) were extracted using the `topTable` function with `coef = “phenotype”`.

#### Functional Enrichment analysis

To identify biological pathways associated with differentially methylated positions, we performed functional enrichment analysis using three complementary approaches. Gene Ontology (GO) enrichment analysis (Biological Process, Molecular Function, and Cellular Component) was conducted using clusterProfiler (v4.0) with gene-centric mapping of CpG sites to their nearest genes. KEGG pathway enrichment was performed using clusterProfiler with the same gene mapping strategy. MSigDB Hallmark gene set enrichment was conducted using missMethyl (v1.26.0), which accounts for the varying number of CpG probes per gene on the Illumina EPIC array. All enrichment analyses were performed independently for each of the 19 facial aging traits using differentially methylated positions at FDR < 0.05 as input. Enriched pathways were considered significant at FDR < 0.05 after Benjamini-Hochberg correction for multiple testing.

#### Interpretation of Effect Sizes

The logFC (log fold-change) coefficient from the limma model represents the estimated change in M-value per unit increase in the phenotype. For continuous HAUT scores (0-100 scale), logFC represents the M-value change per 1-point increase in score (e.g., from score 50 to score 51). For age (years), logFC represents the M-value change per 1-year increase in age. Positive logFC values indicate hypermethylation (increased methylation associated with higher phenotype values), while negative logFC values indicate hypomethylation (decreased methylation associated with higher phenotype values). Because HAUT scores use an inverse convention for many traits (higher scores = better skin quality = less severe phenotype), the biological interpretation of effect direction must account for this scoring system (see Results for detailed interpretation).

#### Multiple Testing Correction

To account for multiple hypothesis testing across 911,580 CpG sites, we applied the Benjamini-Hochberg false discovery rate (FDR) correction using the `adjust.method = “fdr”` option in limma’s `topTable` function. The FDR-adjusted P-values (adj.P.Val) control the expected proportion of false positives among all CpG sites declared significant. We used FDR < 0.05 as the primary significance threshold throughout this study, meaning that among all CpG sites called significant, we expect fewer than 5% to be false positives on average. Secondary significance thresholds of FDR < 0.10 and FDR < 0.01 were also computed for sensitivity analyses.

#### Genomic Inflation Assessment

For each phenotype, we computed the genomic inflation factor (lambda, λ) as the ratio of the median observed chi-squared test statistic to the expected median under the null hypothesis (0.456), based on the conversion of P-values to chi-squared statistics. Lambda values substantially greater than 1.0 can indicate either genuine epigenome-wide signal (many true associations distributed across the genome) or systematic bias (e.g., residual population stratification, uncorrected batch effects, or incomplete adjustment for confounders). We examined quantile-quantile (QQ) plots of observed versus expected P-values for each phenotype to distinguish between these scenarios: genuine polygenic signal manifests as deviation from the null distribution primarily in the upper tail (most significant CpGs), while systematic bias produces uniform inflation across the entire P-value distribution.

#### Cross-Trait Pleiotropy Analysis

To assess the extent of shared epigenetic architecture across skin aging phenotypes, we identified all unique CpG sites reaching significance (FDR < 0.05) in at least one phenotype. For each such CpG site, we counted the number of traits in which it was significant, yielding a “trait count” ranging from 1 (trait-specific) up to 19 (significant in all phenotypes). CpG sites significant in 10 or more traits were classified as “highly pleiotropic” and were examined for evidence of directional consistency (i.e., whether the sign of the effect was consistent across the traits in which the CpG was significant). Pairwise overlap between phenotypes was quantified using the Jaccard similarity index, defined as the size of the intersection (CpGs significant in both traits) divided by the size of the union (CpGs significant in either trait), ranging from 0 (no overlap) to 1 (perfect overlap).

#### Visualization

QQ plots, Manhattan plots, and volcano plots were generated for each phenotype using the ggplot2 package version 3.5.1. Manhattan plots display -log_10_(P-value) for each CpG site on the y-axis against genomic position (chromosome and position) on the x-axis, with horizontal lines indicating genome-wide significance thresholds (FDR < 0.05). Volcano plots display -log_10_(P-value) on the y-axis against logFC on the x-axis, highlighting both statistical significance and magnitude of effect. QQ plots display observed -log_10_(P-values) against expected values under the null hypothesis to assess calibration and identify inflation. All plots were saved as high-resolution JPG images.

#### Software and Package Versions

All analyses were conducted in R version 4.4.0 (2024-04-24, “Puppy Cup”). Key R packages and versions used include:

1. *Methylation data processing:*
  a. *SeSAMe (version 1*.*20*.*0): Preprocessing, quality control, normalization (ssNoob method)*
  b. *IlluminaHumanMethylationEPICv2manifest (version 0*.*99*.*0): Array manifest*
  c. *IlluminaHumanMethylationEPICv2anno*.*20a1*.*hg38 (version 0*.*99*.*0): CpG annotation*
2. *Cell type deconvolution:*
  a. *EpiSCORE (version 1*.*0*.*0): Reference-based deconvolution with mrefSkin*.*m*
3. *Differential methylation analysis:*
  a. *limma (version 3*.*58*.*1): Linear models and empirical Bayes moderation*
  b. *DMRcate (version 3*.*0*.*0): Differentially methylated region identification (planned for future analysis)*
4. *Data manipulation and visualization:*
  a. *data*.*table (version 1*.*15*.*4): Efficient data import and manipulation*
  b. *dplyr (version 1*.*1*.*4): Data wrangling*
  c. *tidyr (version 1*.*3*.*1): Data tidying*
  d. *ggplot2 (version 3*.*5*.*1): Publication-quality graphicsggrepel (version 0*.*9*.*5): Non-overlapping text labels in plots*

Scripts implementing these analyses are available from the authors upon request.

## Acknowledgements

We would like to thank Chatlin Helm and Razelle Ann Piculados for their valuable help with data management. We would also like to thank the many volunteers who supported this study by providing skin samples which were used for the development of predictors.

## Data Availability

Due to privacy considerations and institutional review board restrictions, individual-level methylation and phenotype data cannot be publicly shared. The HAUT Face Analysis 3.0 software is proprietary and available under commercial license from Haut.AI (https://haut.ai). Raw methylation array data (Illumina EPIC v2) will be available upon reasonable request. The crosswalk linking anonymous subject identifiers to internal sample IDs is available from the corresponding author under a data use agreement (DUA) to protect participant privacy. Additional data supporting the findings of this study are available from the corresponding author upon reasonable request. Full supplementary data and information is available on Zenodo at https://zenodo.org/records/19006544.

## Code Availability

Analysis code for the EWAS, cross-trait pleiotropy analysis, lifestyle association analyses, and figure generation will be deposited on Zenodo (https://zenodo.org/records/19006544) upon acceptance and made publicly available at that time.

## Author Contributions

Design and conceptualization: VBD, RS [Ryan Smith], LP. Methylation data generation and processing: VBD. Statistical analyses: VBD, SH. Predictive model development: SH, VBD. Clinical data generation and sample provision: RS [Sivamani], JG. Facial imaging data generation and processing: AG, KK. Sample collection and processing: DS, JW. Scientific discussion and interpretation: GMF, YC, KS. Manuscript drafting: VBD, SH. All authors reviewed, edited, and approved the final manuscript.

## Competing Interests

VBD, SH, KS, DS, JW, and RMS are employees of TruDiagnostic Inc. AG and KK are employees of Haut.AI. LP is an employee of Olaplex Inc. GMF, YC, JG, and RS declare no competing interests.

## Consent for Publication

Facial images were collected for automated phenotyping via HAUT.ai but are not reproduced in this article. All participants provided informed consent for the use of their biological samples and imaging data in research. No identifiable participant data appear in this publication.

## Supplementary Figure

**Supplementary FIGURE S1.**
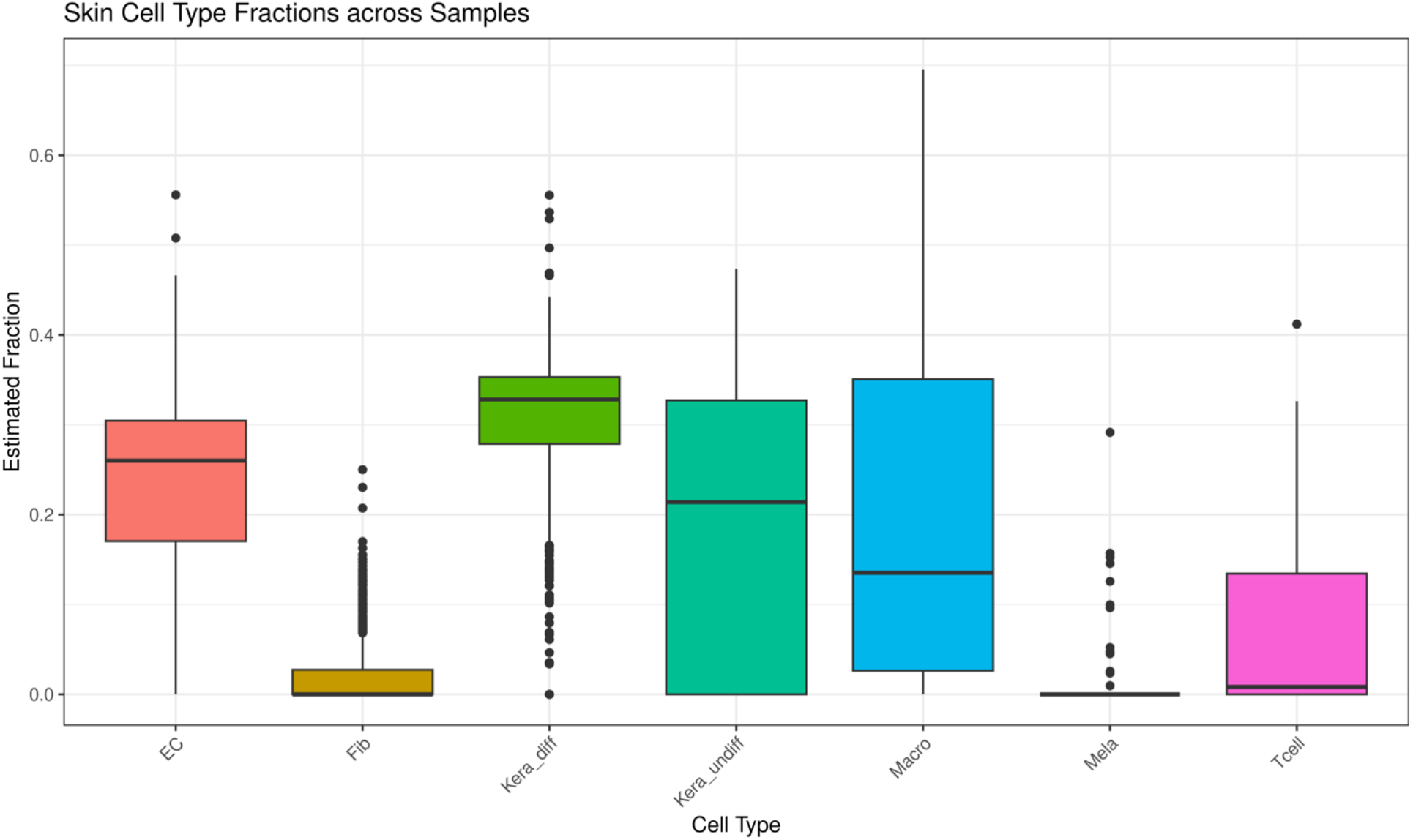
Skin cell-type composition estimates across samples. Boxplots summarize EpiSCORE-inferred skin cell-type fractions across 744 facial skin samples, including endothelial cells (EC), fibroblasts (Fib), differentiated keratinocytes (Kera_diff), undifferentiated keratinocytes (Kera_undiff), macrophages (Macro), melanocytes (Mela), and T cells (Tcell). Fractions are expressed as percentages summing to ∼100% per sample. Differentiated keratinocytes comprised the largest component (median 32.82%), followed by endothelial cells (26.01%) and undifferentiated keratinocytes (21.39%). Macrophage fractions were variable (median 13.53%), whereas T cells were lower on average (median 0.83%). Fibroblast (median 0.00%) and melanocyte (median 0.00%) fractions were generally minimal across samples. Center lines indicate the median, boxes denote the interquartile range (Q1–Q3), whiskers extend to 1.5×IQR, and points represent outliers. The predominance of keratinocytes with a notable endothelial contribution is consistent with superficial, epidermis-enriched sampling, and these estimates were included as covariates in downstream methylation analyses to account for cellular heterogeneity.

## Supplementary File Legends

**SUPPLEMENTARY FILE 1** | **Cohort characteristics and demographic composition**. Summary of sample collection, data availability, and key demographics for the HAUT imaging cohort, the Illumina EPIC v2 methylation cohort, the survey-linked subset, and the matched HAUT–methylation subset used for model development and downstream analyses. Age statistics are reported for participants with available metadata. Fitzpatrick skin type counts are shown for the methylation cohort; “not reported” indicates missing Fitzpatrick data. ^a^Sample size varies across the 21 EpiVision predictors due to phenotype-specific completeness and covariate availability.

**SUPPLEMENTARY FILE 2** | **Complete performance metrics for all EPICv2 (V2-Only) and V1–V2 overlap predictive models across skin trait targets**. All metrics are reported on the held-out test set (20% of total samples). ΔMAE and ΔSpearman R reflect the percent change of the EPICv2 model relative to the V1–V2 overlap model (negative ΔMAE indicates improvement; positive ΔSpearman R indicates improvement). Targets are sorted alphabetically. Probe counts: EPICv2 probe set uses all stable EPICv2 probes; V1–V2 overlap probe set is restricted to probes present on both EPICv1 and EPICv2 arrays.

**SUPPLEMENTARY FILE 3** | **Association results between self-reported lifestyle, clinical, and medication exposures and EpiVision methylation-based facial aging predictors**. Results are reported for 33 exposures tested against 21 EpiVision predictors in the survey-linked subset (n = 282) across four models: unadjusted raw predicted age (Age – Raw); epigenetic age acceleration (Age – EAA), in which predicted age is residualized on chronological age to isolate biological aging effects independent of time; unadjusted phenotype scores (Phenotype – Raw); and age/sex-adjusted phenotype scores (Phenotype – Model 2), restricted to participants ≥16 years. Exposures include binary clinical and medication variables (e.g., retinoid use, hormone therapy, spironolactone, testosterone) and ordinal lifestyle variables (e.g., sleep quality, diet quality, water intake, tobacco status). Standardized beta coefficients are reported throughout to enable cross-exposure comparability. FDR correction applied within adjusted models using the Benjamini– Hochberg method. Associations highlighted in the main text reach p < 0.05 in Model 2 with n_exposed > 1; associations with n_exposed = 1 are considered exploratory.

**SUPPLEMENTARY FILE 4** | **Paired comparison of HAUT-derived skin phenotype scores between sun-protected and sun-exposed skin sites**. Results from paired t-tests comparing 21 facial aging and skin quality phenotypes between sun-protected buttock skin and sun-exposed forearm skin collected from the same individuals (n = 20 per trait). Traits are sorted by ascending p-value. Mean Difference reflects forearm minus buttock values; negative values indicate lower scores in sun-exposed skin (i.e., worse phenotype given HAUT’s scoring convention where higher = better quality), and positive values indicate higher scores in sun-exposed skin. Cohen’s d_z is the within-subject standardized mean difference, calculated as the mean paired difference divided by the standard deviation of the paired differences, and reflects effect size for paired designs. The Panel column corresponds to figure panel labels in the associated supplementary figure. Statistical significance threshold: p < 0.05 (two-sided paired t-test, no correction for multiple comparisons across traits).

**SUPPLEMENTARY FILE 5** | **Epigenome-Wide Association Study Results for 19 Skin Aging and Appearance Traits**. N, sample size; Lambda, genomic inflation factor; DMPs, differentially methylated positions at FDR < 0.05; % Sig, percentage of tested CpGs (911,580 total) reaching significance; % Hyper, percentage of significant DMPs showing hypermethylation (positive effect); Top CpG, most significant DMP by P-value; logFC, log fold-change in M-values per unit increase in phenotype. All models adjusted for technical batch (Illumina BeadChip) and six skin cell type fractions (fibroblasts, differentiated keratinocytes, undifferentiated keratinocytes, macrophages, melanocytes, T cells) estimated using EpiSCORE with skin-specific reference.

**SUPPLEMENTARY FILE 6** | **Functional enrichment analyses for differentially methylated positions associated with 19 facial aging and skin quality traits**. Pathway enrichment was performed independently for each trait using genes mapped to significant DMPs (FDR < 0.05) identified in the covariate-adjusted EWAS, across four complementary databases: Gene Ontology Biological Process (GO_BP), Molecular Function (GO_MF), and Cellular Component (GO_CC); KEGG Pathways; and MSigDB Hallmark gene sets. In total, 28,583 enriched pathways were identified across all 19 traits, sorted by trait and ascending p-value. Significance threshold: FDR < 0.05.

