## Supplementary figures and images for "Skin DNA Methylation Encodes Multidimensional Facial Aging Phenotypes with Distinct Biological Architectures"

### Figure1.tiff

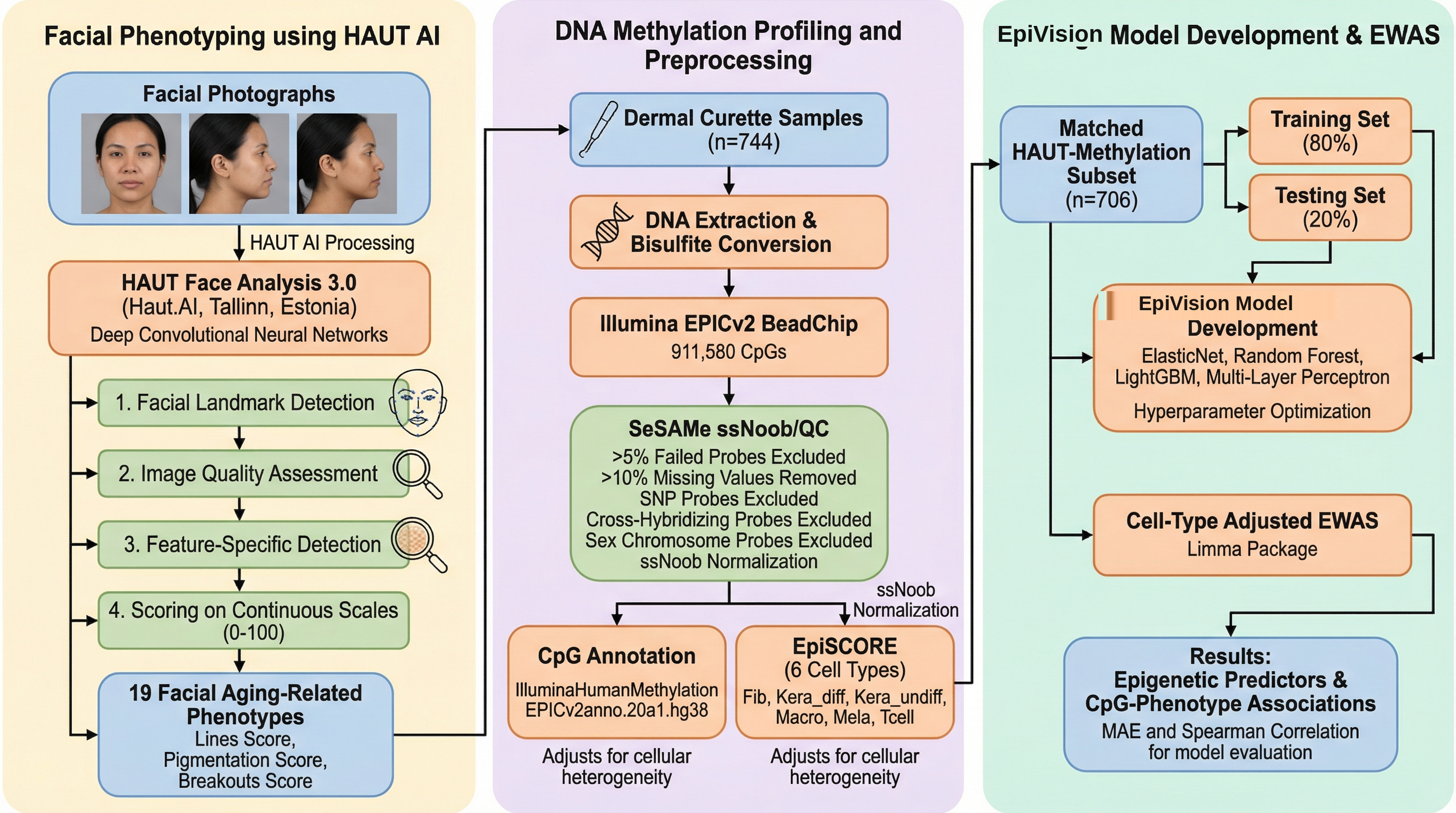

### Figure2.tiff

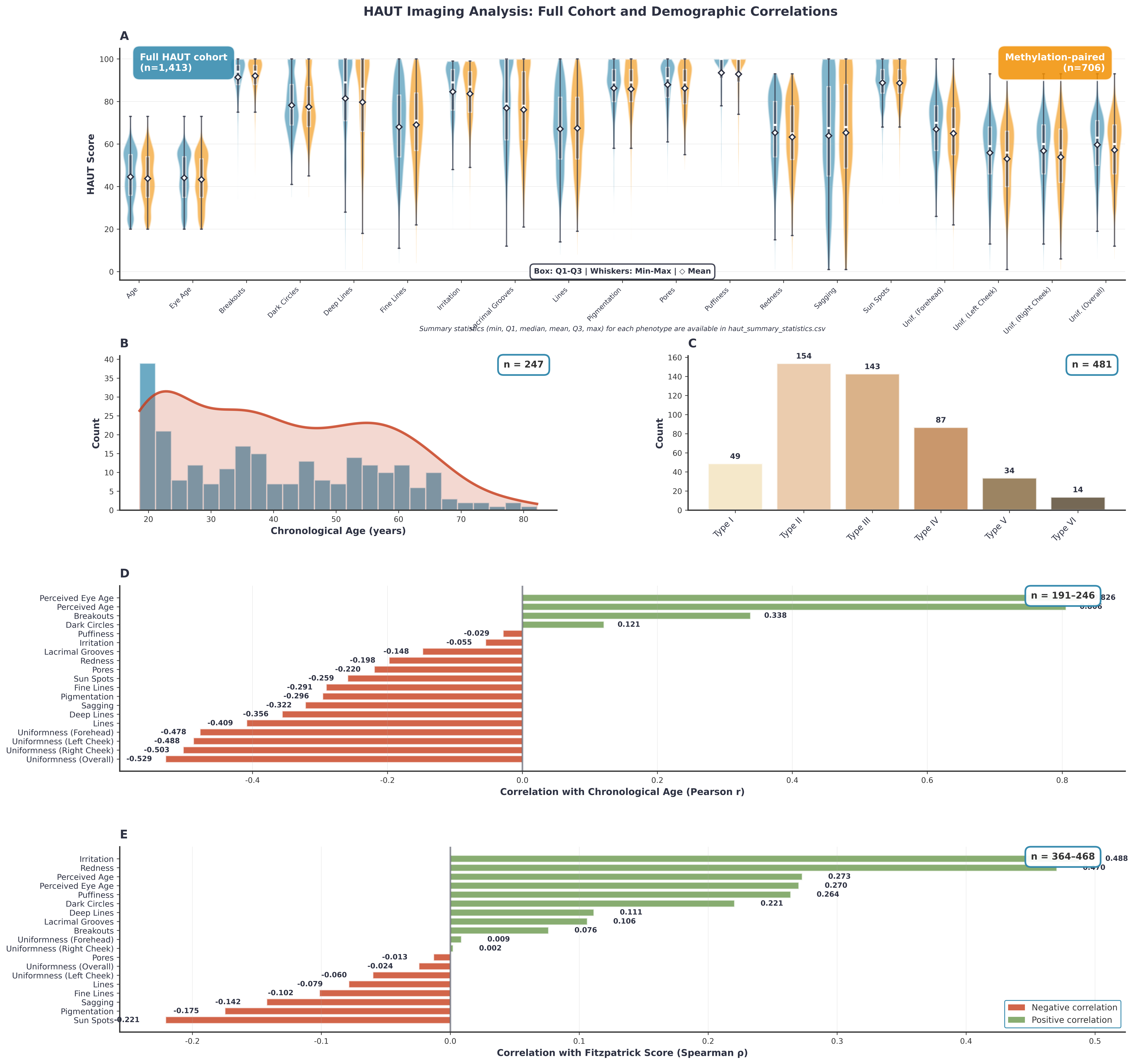

### Figure3.tiff

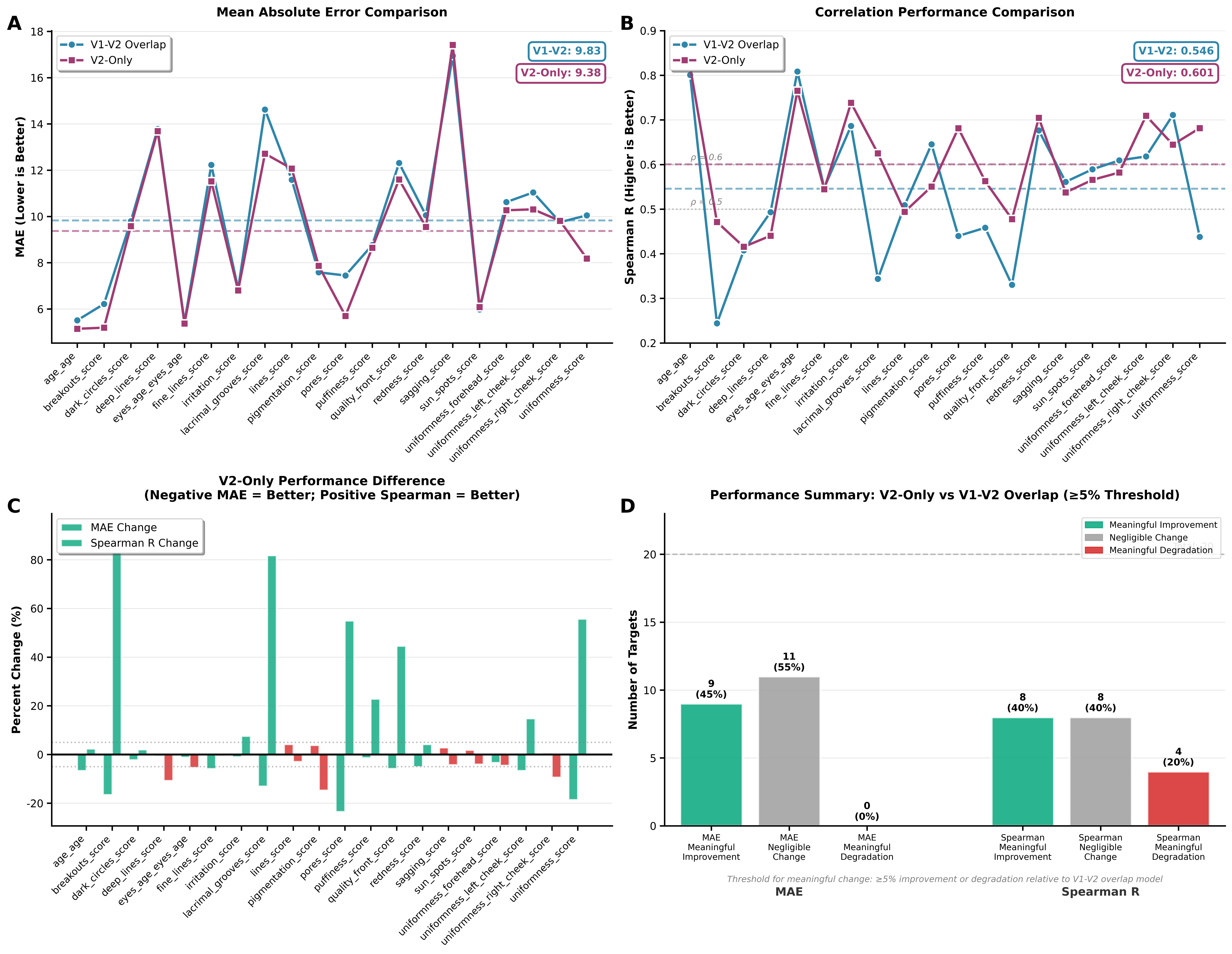

### Figure4.tiff

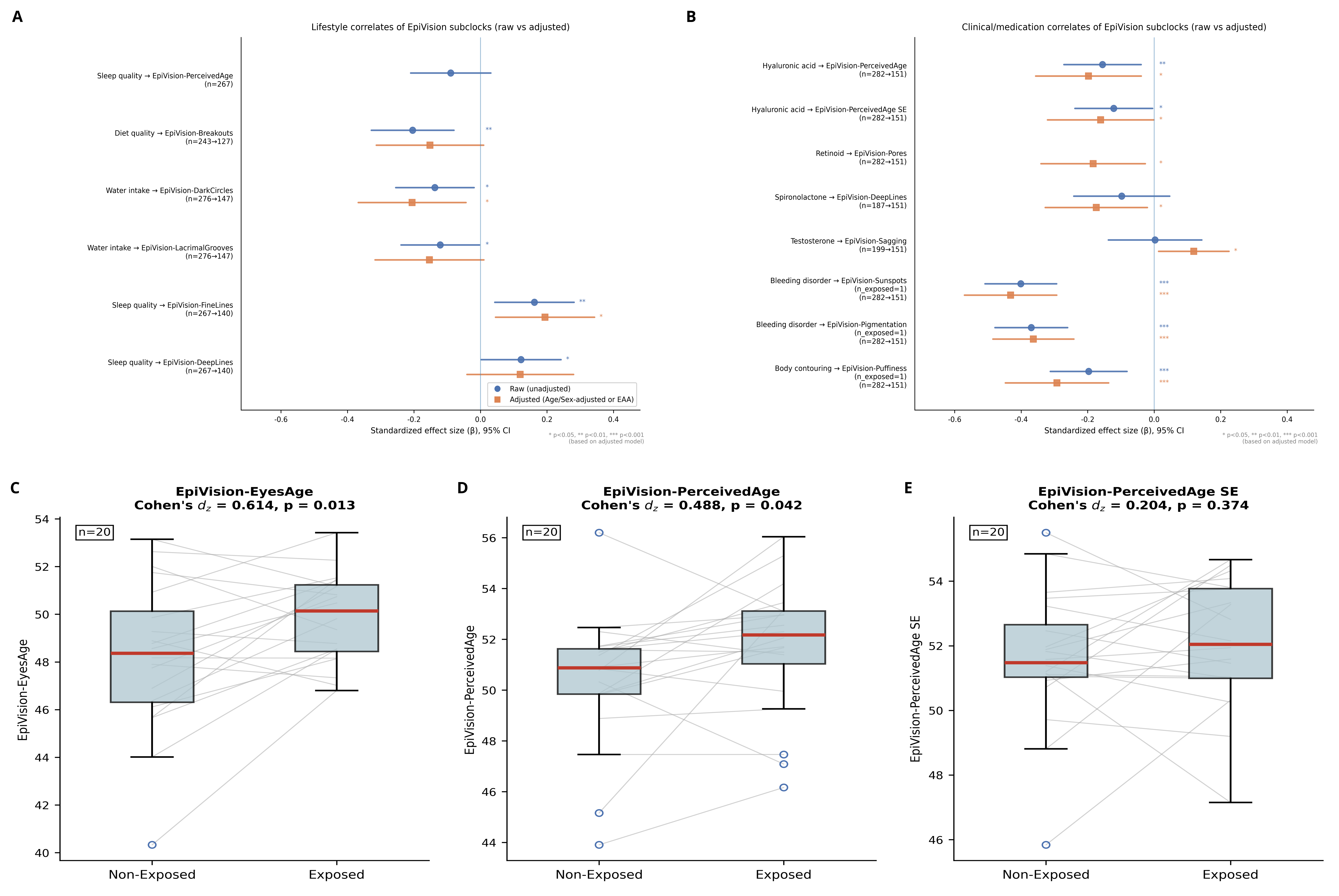

### FigureS1 (1).png

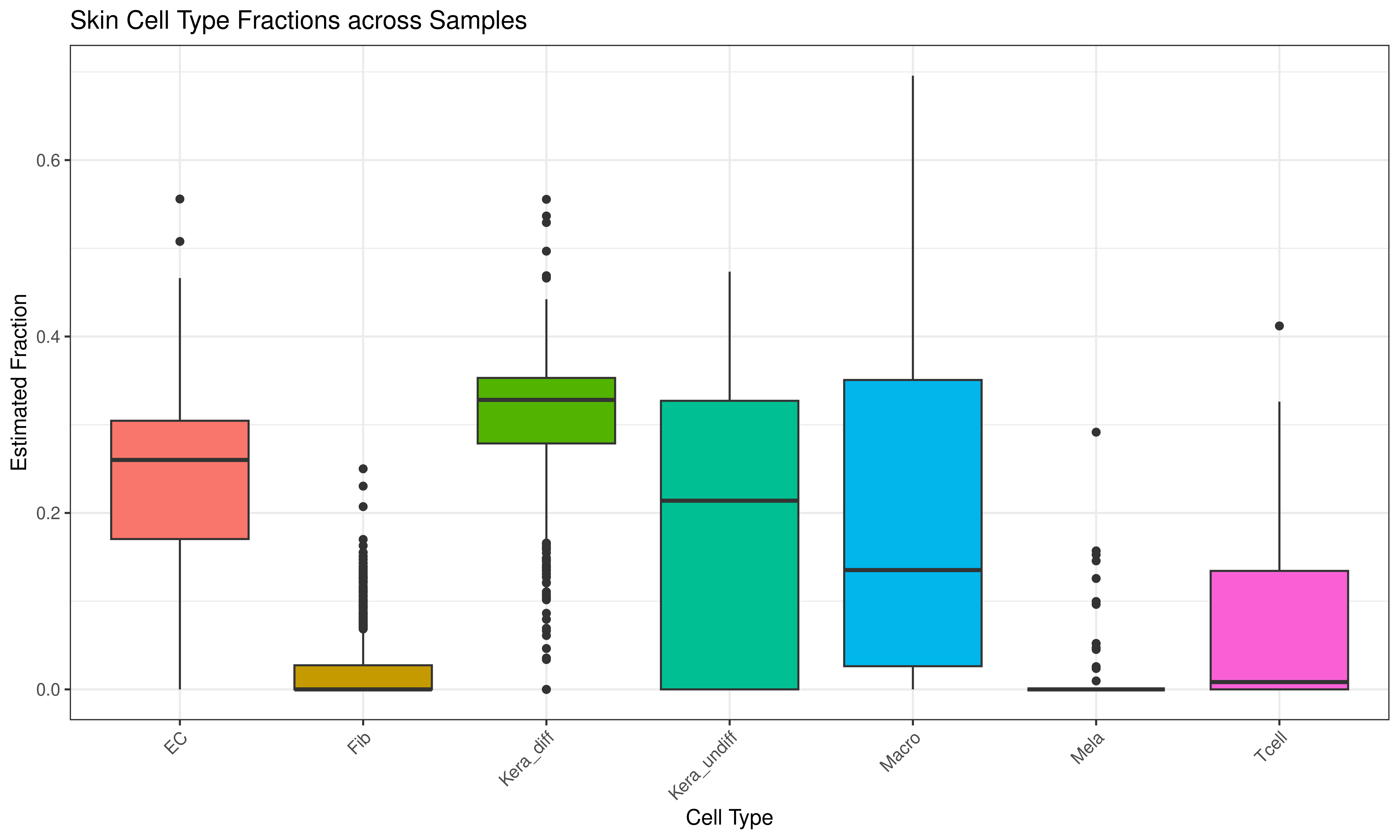
